# Electrical Automaticity and Intercellular Synchronization via Shared Extracellular Compartments

**DOI:** 10.1101/2020.04.15.043414

**Authors:** St. Poelzing, J. P Keener

**Affiliations:** Virginia Tech Carilion; University of Utah

## Abstract

Electrically excitable cells often spontaneously and synchronously depolarize in vitro and in vivo. It remains unknown how cells synchronize and autorhythmically activate above the intrinsic mean activation frequency of isolated cells without pacemaking mechanisms. Recent insights into ephaptic coupling (non-gap junction or synaptic coupling) suggest that cyclic ion accumulation and depletion in diffusion limited extracellular volumes densely expressing ion channels modifies action potentials. This report explores how potassium accumulation and depletion in a restricted extracellular domain promotes spontaneous oscillations in the Hodgkin Huxley action potential model, which does not spontaneously activate on its own without external stimulus. Simulations demonstrate cells sharing a diffusion limited extracellular compartment can become autorhythmic and synchronous despite intercellular electrical heterogeneity. Autorhythmic frequency can be determined by net potassium flux into the cleft and the cleft volume. Additionally, inexcitable cells can induce autorhythmic activity in an excitable cell via a shared cleft and sufficient potassium fluxes contributed by each cell. Importantly, the synchronization and autorhythmic activity conferred by shared cleft with reduced potassium efflux can occur in the absence of gap junctions. Lastly, not only can potassium oscillations in shared restricted clefts initiate, support, and suppress autorhythmic depolarizations, the same mechanism can homogenize repolarization. The work has implications for understanding how automaticity is coordinated among excitable cells and suggests a new role for non-excitable cells such as fibroblasts, macrophages, or astrocytes with sarcolemmal potassium handling proteins facing shared and restricted intercellular clefts.

**SIGNIFICANCE:** A mechanism of cyclic ion accumulation and depletion in diffusion limited extracellular compartments can suppress, initiate, and support autorhythmic activity. Additionally, autorhythmicity can emerge from electrophysiologically heterogeneous cell pairs sharing a diffusion limited extracellular compartment, even if the individual cells will not spontaneously depolarize on their own. Sustained and synchronous autorhythmic activity can occur in the absence of gap junction coupling. Lastly, the shared diffusion limited extracellular compartment can also reduce action potential duration gradients by synchronizing repolarization.

## INTRODUCTION

Cyclic behavior is frequently observed in multicellular preparations. At the cellular level, oscillations often occur secondary to linked processes with different reaction time constants, positive and negative feedback, and reagent availability. Oscillators can synchronize or continue to function independently with minimal energy or resource transfer. In the context of excitable cells physically defined and constrained by a cell membrane such as neurons, or myocytes, cyclic electrical activity arises from regenerative sarcolemmal ion exchange or other intracellular ionically dependent processes. For example, there are two predominant theories that describe the origin of the heartbeat in specialized cells within the sinoatrial node. It is well accepted that the “funny current", linked to the hyperpolarization-activated, cyclic-nucleotide gated family of genes (HCN), is a steady-state pacemaking current that predominates in late diastole, permitting slow membrane depolarization that eventually triggers an action potential AP (1)(2)(3)(4). According to oscillatory systems theory, some type of feedback that can be positive or negative is required to reset, limit, or select for a dominant frequency. In the case of the cardiac myocyte membrane potential, this is accomplished by the voltage and time dependent activation, inactivation and deactivation of a host of ion handling proteins known as channels, exchangers and pumps. The setting of intrinsic cyclic depolarization is referred to as the “membrane clock” hypothesis, and the structure, function and modulation of the HCN channels were elegantly summarized by Wilders in 2007 (5).

The “calcium clock” hypothesis, which can operate in conjunction with the membrane clock, was reviewed carefully by Lakatta and colleagues in 2010 (6). In short, the calcium clock operates with the assumptions that calcium regulatory channel proteins called ryanodine receptors on the sarcoplasmic reticulum will spontaneously and nearly synchronously release calcium into the cytosol. Some of that calcium will leave the cell through a sodium-calcium exchanger. The exchange of 1 calcium for 3 sodium ions depolarizes the myocyte by a net charge transfer, providing the positive and negative feedback for additional calcium entry and exchange through L-type calcium channels, further raising intracellular calcium. Some of this calcium leaves the cell again, but the majority of the calcium is taken back into the sarcoplasmic reticulum through a pump. As the system equilibrates between action potentials, increased SR calcium increases the likelihood of spontaneous calcium release again.

Potassium handling has also been implicated as an important mechanism of electrical automaticity in cardiomyocytes. Specifically, many excitable cells have a negative resting membrane potential, and this is generally attributed to inwardly rectifying potassium channels that polarize the membrane closer to the Nernst reversal equilibrium potential of potassium. The role of potassium channels and many other sarcolemmal currents are summarized by Irasawa et al.(7) In brief, the inward rectifier potassium channels are thought to inhibit automaticity, and therefore altering potassium permeability is a second membrane clock mechanism that can drive intrinsic cyclic behavior. In cardiac electrophysiology, this is what is thought to underlie the different and increasingly slower spontaneous action potential rates distal to the sinoatrial node in the atrioventricular node, His bundle, Purkinje fibers, and eventually the working myocardium. Beyond the heart, spontaneous and cyclic electrical behavior has been described in the brain (8)(9)(10), hair cells (11), skeletal myotubes (12) and uterine myometrium (13). In short, excitable tissues in general are associated with varying degrees of coordinated cyclic electrophysiologic action potential activity.

However, a consensus on what modulates and regulates organized cyclic activity in a group of cells is less well developed. Consider that the probability of a group of cardiac cells depolarizing sufficiently to initiate coordinated and near simultaneous depolarization is proportional to the probability of a single cell depolarizing at any given point in time raised to the power of the number of other cells that also need to spontaneously and nearly synchronously depolarize. With this thought experiment, this means that even if an individual cell has a 90% probability of spontaneously activating at a given time, the probability of 100 cells depolarizing at the same time is extremely small, expressed as 0.90^100^ = 2.7 × 10^−5^. Some have even estimated it would take 10,000 cardiomyocytes to simultaneously depolarize and provide sufficient excitatory current to propagate and cause spontaneous contraction of cardiac tissue by this mechanism (14).

In short, following relatively straightforward mathematical assumptions, the likelihood of synchronized electrical activity in groups of cells is unlikely without a pacemaking mechanism. As a result, the fundamental mechanisms underlying cellular synchronization remain elusive, and this gap in knowledge is relevant in the investigation of epilepsy and sudden cardiac death. Many theories have been proposed utilizing advanced mathematics to describe how synchronized, entrained, and resonant activity can emergently arise in tissue, and this study does not challenge any previous work. Rather, this report describes the logic and a simplified Hodgkin-Huxley (HH) mathematical action potential model (15) showing how a single cell with no intrinsic oscillatory activity can gain oscillatory activity and even accelerate spontaneous activity in a group of cells coupled via shared diffusion limited extracellular compartments. Importantly, this theory does not require gap junctions or synapses and overcomes the issues concerning stochastic behavior in large populations. The model is based on recent findings from multiple investigators that the intercalated disc of cardiac myocytes densely express sodium and potassium channels (16)–(26). The description of the gap junction adjacent perinexus, first reported by Rhett, Jourdan, and Gourdie in 2011 (27), lead to our proposition that altering the size of this nanodomain can conceal a sodium channel gain of function pathology by a mechanism of sodium ion depletion in the intercalated disc (28)(29). Briefly, narrow perinexi support a transient state where sodium channels are highly conductive but will not pass current because the driving force - the difference between the membrane potential (*V*_*m*_) and sodium reversal potential (*E*_*Na*_) - is substantially reduced towards zero.

## METHODS

The first significant assumption of our model is based on studies demonstrating that a subpopulation of inward rectifier potassium channels (Kir2.1) are localized to the relatively restricted volume of the ventricular myocardial intercalated disc (25), which is referred to as the “cleft” in this model. The second assumption is that extracellular potassium concentration in the cleft can vary more than, and independently of, intracellular and bulk extracellular potassium. For context, the extracellular volume in working ventricular myocardium is tens of microns wide and spatially extensive throughout tissue, intracellular dimensions of the adult human ventricular cardiomyocyte are approximately 20 by 100*μ*m (30), and the perinexus is approximately 20nm wide and approximately 100nm long (24)(31). Thus, the perinexus is several orders of magnitude smaller in volume than cardiomyocytes and the interstitial volume.

Given these assumptions, the HH model of a single cell was altered according to Figure 1. Specifically, membrane currents were considered to consist of two components, the lateral membrane with membrane area *A*_*m*_ and the junctional membrane, i.e., the membrane lining the cleft with membrane area *A*_*j*_. Each of the membrane currents consist of capacitive currents and ionic currents with sodium, potassium and leak ionic currents for the lateral membrane and sodium and potassium ionic currents for the junctional membrane. Consequently, the total transmembrane current is

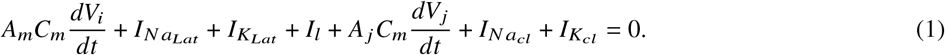

**Figure 1:**
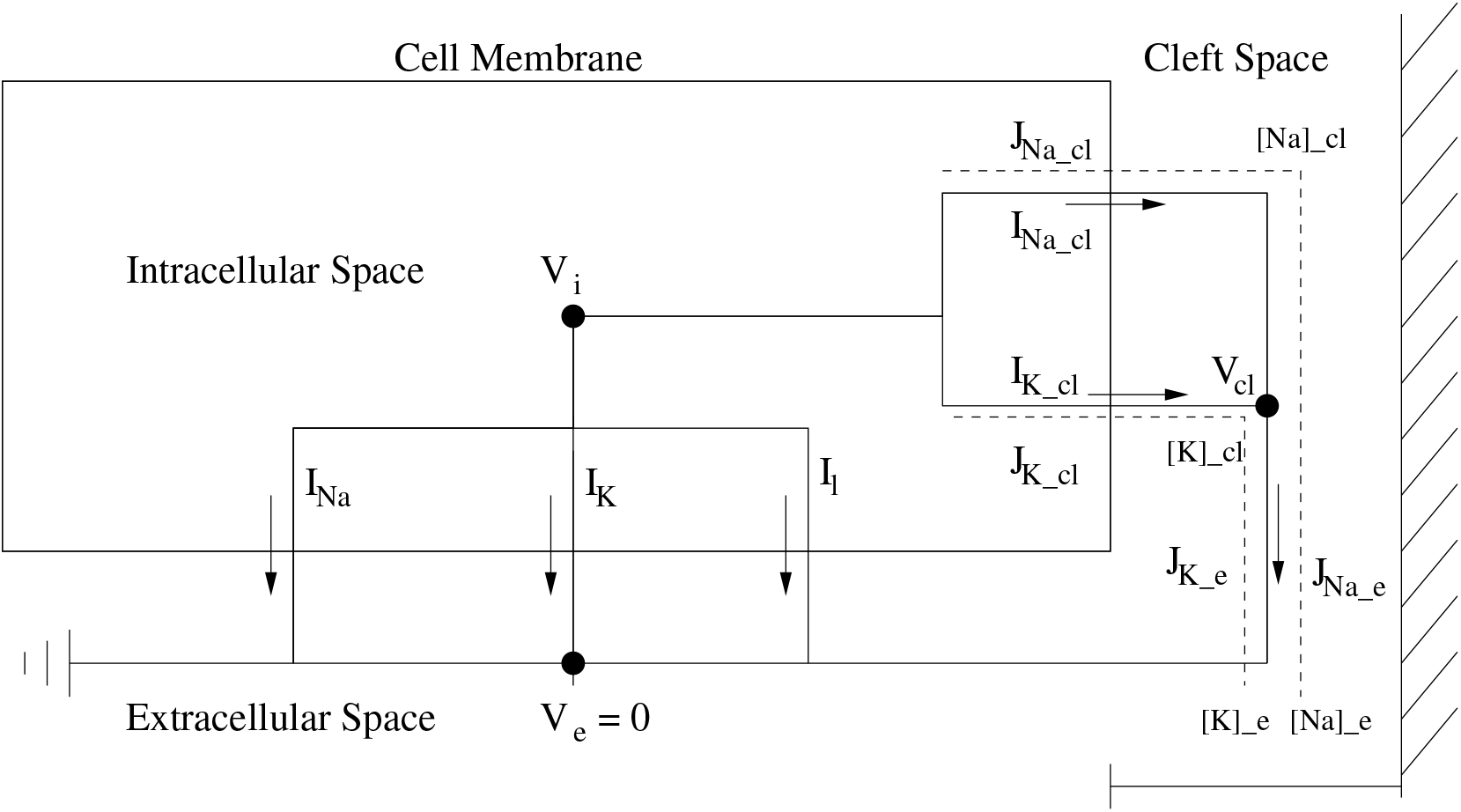
Circuit diagram for a single Hodgkin-Huxley cell modified to include sodium and potassium currents into a narrow cleft region. Model includes standard HH transmembrane ionic currents for the lateral membrane and potassium and sodium transmembrane currents for the junctional membrane into the narrow cleft region. Currents out of the cleft region are governed by a GHK current relationship. The two cell model has a single and shared [*K*]_*cl*_ (cleft potassium concentration) and [*Na*]_*cl*_ (cleft sodium concentration), *w* _*f*_ (cleft width), and *r*_*j*_ (cleft resistance) parameters. Cellular capacitance, and fraction (*F*_*I K*_) of total potassium current 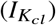 into the cleft can be varied independently for both cells. Supplemental files single_cell_potassium_cleft.m and double_cell_potassium_cleft.m are Matlab source code for the single and dual cell models, respectively.

The lateral membrane has transmembrane potential *V*_*m*_ = *V*_*i*_ − *V*_*e*_ (with *V*_*e*_ = 0) and the junctional membrane has transmembrane potential *V*_*j*_ = *V*_*i*_ − *V*_*cl*_, where *V*_*cl*_ is the potential in the cleft and is not the same as *V*_*e*_. The junctional membrane current must balance the cleft current *I*_*cl*_,

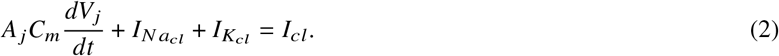

The lateral membrane ionic currents are gated in a voltage-dependent way by *V*_*i*_, whereas the junctional membrane ionic currents are gated in a voltage-dependent way by *V*_*j*_. Furthermore, the driving force (the Nernst potentials) for the lateral membrane ionic currents are fixed at standard values, however, the Nernst potentials for the junctional ionic currents fluctuate with the cleft concentrations according to

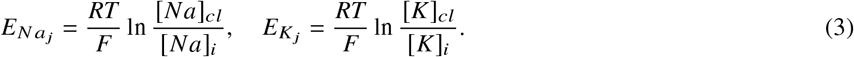

Because the cleft volume is so small, we track the cleft potassium and cleft sodium concentrations by

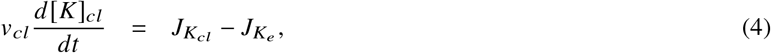

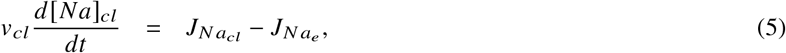

where the cleft volume is *v*_*cl*_ = *A*_*j*_*w*, *w* is the cleft width. Ionic fluxes *J* and currents *I* are related through Faraday’s constant

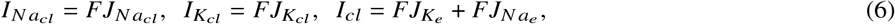

Finally, for the cleft currents and ion fluxes, we take the Goldman-Hodgkin-Katz (GHK) fluxes

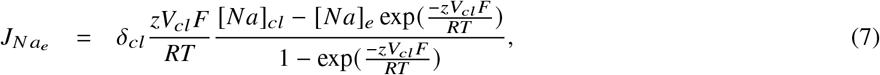

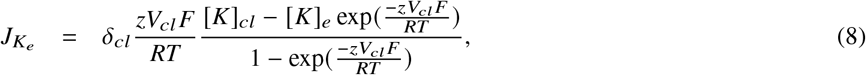

with *z* = 1. Notice that if *V*_*cl*_ = 0, these reduce to Fickian diffusion, whereas with no concentration gradient, these reduce to

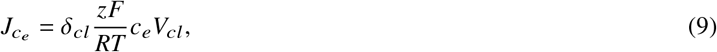

which is Ohm’s law. Thus, since there are two ions involved, we take

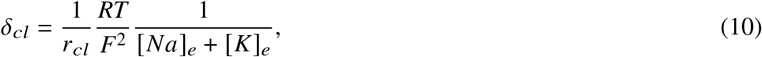

where *r*_*cl*_ is the cleft resistance.

The model has three adjustable parameters. These are *F*_*I K*_, *F*_*I N a*_, and *w* _*f*_. *F*_*I K*_ and *F*_*I N a*_ specify the fraction of potassium and sodium ion channels, respectively, in the junctional membrane, and are between zero and one. For all the simulations presented here, *F*_*I N a*_ = 0, so that the junctional membrane has no sodium ion current. The cleft width factor *w* _*f*_ appears in two places, in the cleft volume *v*_*cl*_ = *A*_*j*_ *w* _*f*_ *w*^0^, where *w*^0^ is the nominal cleft width, and in the cleft resistance 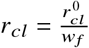, where 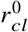 is the nominal cleft resistance. Consequently, the cleft diffusion coefficient is 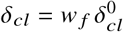.

Parameter values used for this study are listed in Table 1.

**Table 1:**
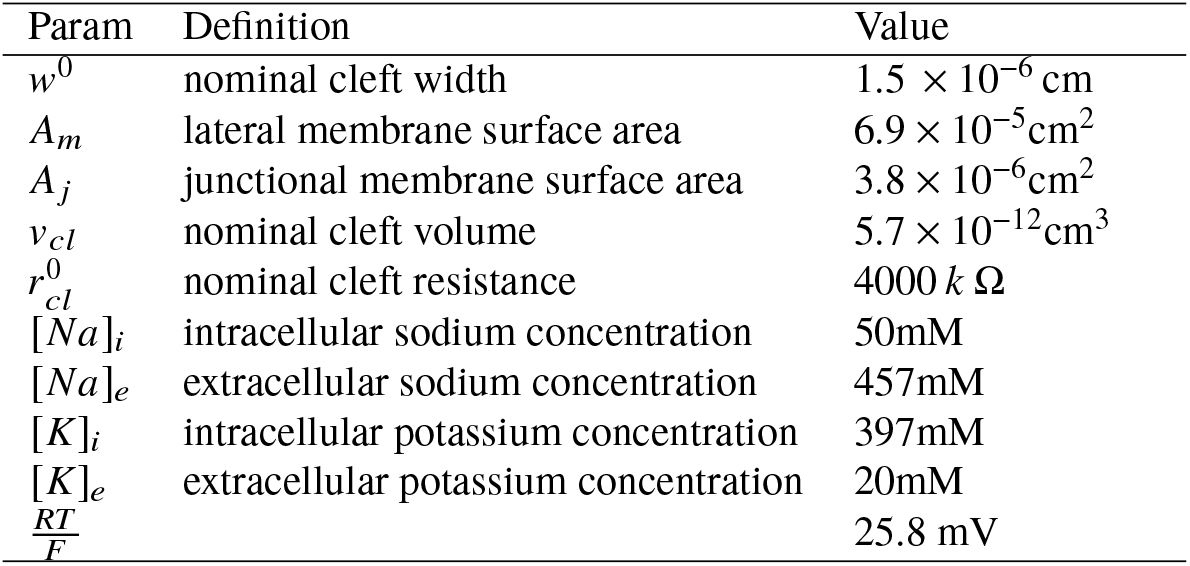
The definition and value of parameters

## RESULTS

### Oscillatory Behavior with Potassium Diffusion Limited Clefts

The HH model is not spontaneously oscillatory with standard parameter values without the addition of a transient or persistent depolarizing current. However, it does exhibit oscillatory behavior if the extracellular potassium concentration is elevated. This fact is demonstrated in Fig. 2. In panel A, the red curves represent stable steady solutions, and the blue curve represents stable oscillatory solutions. Consequently, for 29mM < [*K*]_*e*_ < 58mM, there are stable oscillatory solutions. In Fig. 2B, the oscillatory solution is shown for [*K*]_*e*_ = 30mM. The mechanism underlying these oscillations is easy to understand. The increased extracellular potassium concentration results in an increase in the potassium Nernst potential which effectively provides a constant depolarizing current, relative to the standard value at [*K*]_*e*_ = 20mM, and oscillations result.

**Figure 2:**
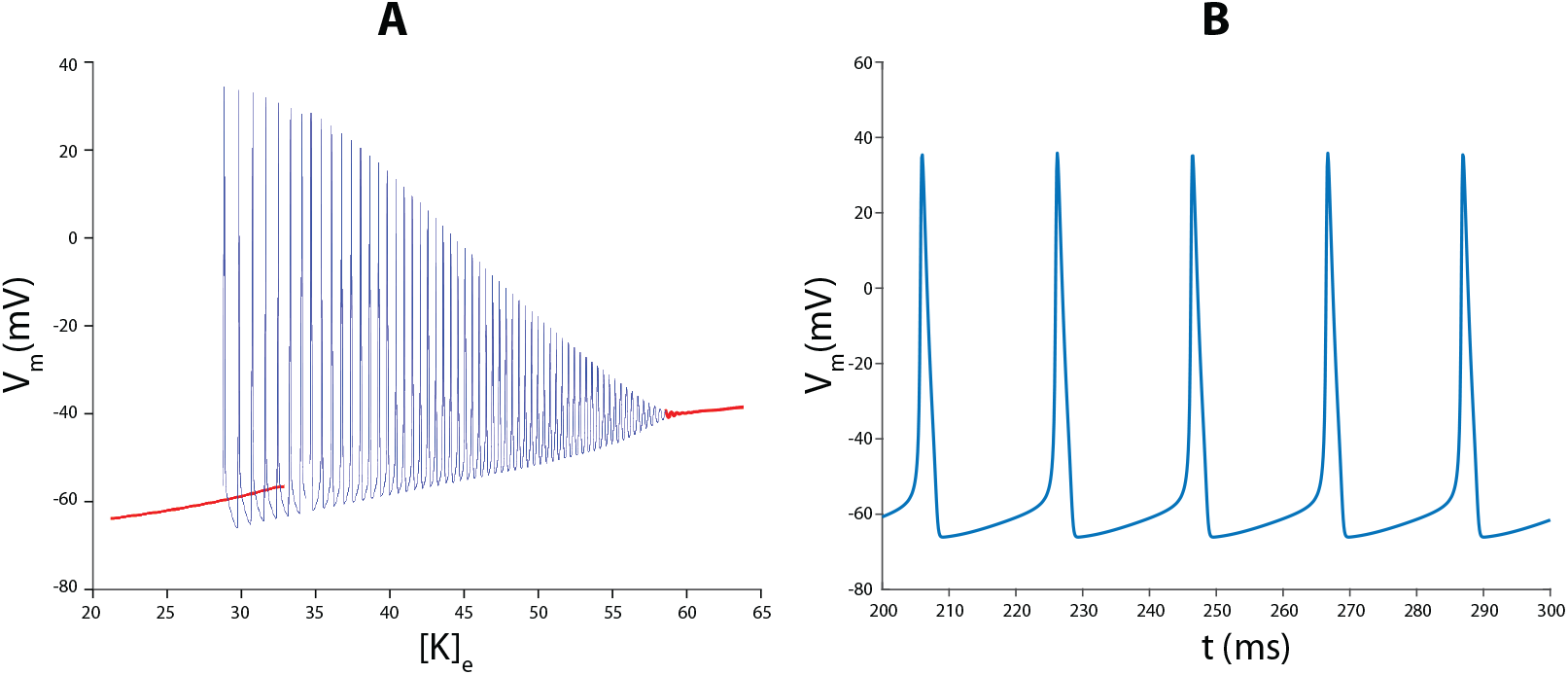
Panel A: Solution diagram for the HH model as a function of extracellular potassium concentration [*K*]_*e*_. Red curves represent stable steady state solutions and the blue curve represents stable oscillatory solutions. Panel B: Spontaneous oscillatory solution for the HH model with [*K*]_*e*_ = 30mM.

To explore whether spontaneous activity can be initiated by extracellular potassium control in our modified model, the total potassium current *I*_*K*_ was distributed between channels facing an invariant potassium compartment and the cleft where potassium concentration can change. Figure 3 demonstrates that with the fraction of potassium channels facing the cleft, *F*_*I K*_ = 0.7, there is spontaneous oscillatory behavior.

**Figure 3:**
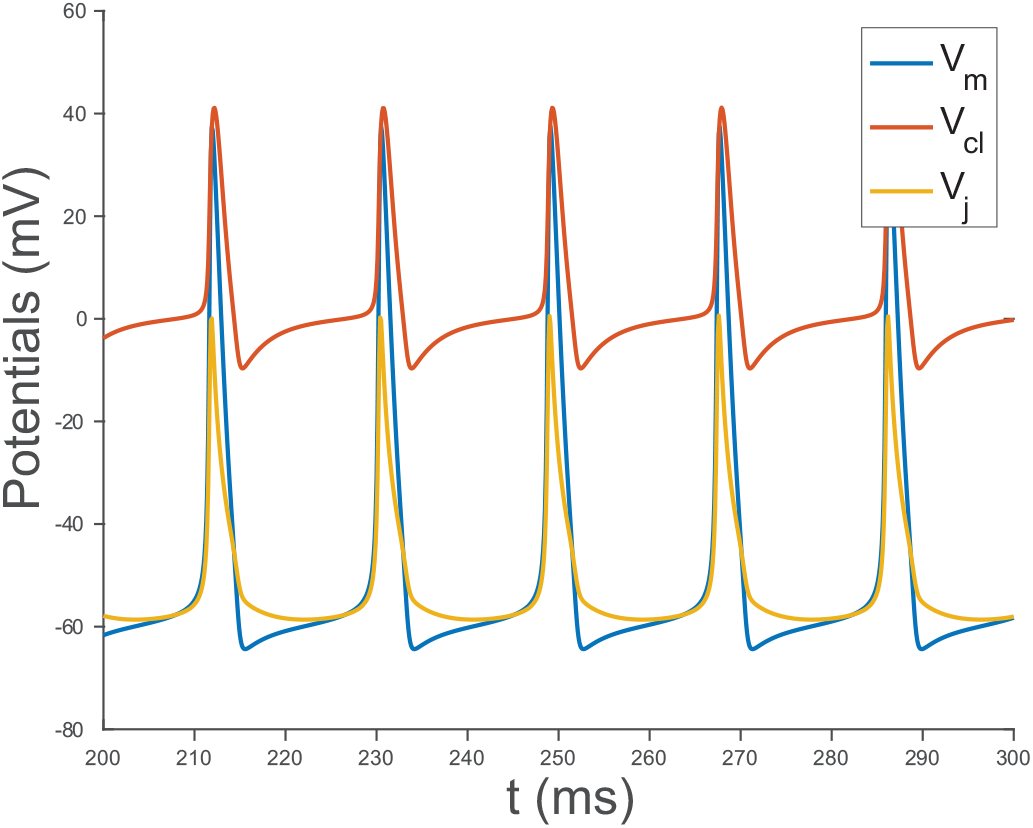
Spontaneous oscillatory behavior for the membrane potential *V*_*m*_ = *V*_*i*_, the cleft potential *V*_*cl*_ and the junctional transmembrane potential *V*_*j*_ = *V*_*i*_ − *V*_*cl*_ with fraction of potassium cleft current *F*_*I K*_ = 0.7 and nominal cleft width *w* _*f*_ = 1.

The mechanism behind the spontaneous activity is illustrated in Figure 4. First, the lateral membrane potential (*V*_*m*_) does not drop below *E*_*K*_, because *E*_*K*_ = −65 mV is the most negative reversal potential for the bulk membrane. As a result, potassium current along the lateral membrane 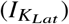 facing an unrestricted volume can only ever be outward (Panel A). Panel B demonstrates that facilitating a change in [*K*]_*cl*_, resulting from 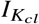, permits transient cyclic [*K*]_*cl*_ increases. In the cleft, [*K*]_*cl*_ does not drop below 40mM, which is higher than the lateral and invariant potassium concentration of 20mM. The consequence is that the junctional Nernst potential 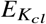 can transiently and cyclically rise above resting *V*_*m*_ (Panel C). When 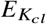 rises above *V*_*m*_, then 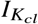 reverses to become an inward current (Panel D). It is this mechanism of inward potassium current caused by extracellular potassium accumulation in the cleft that causes periodic oscillations of the HH model without the injection of any additional current.

**Figure 4:**
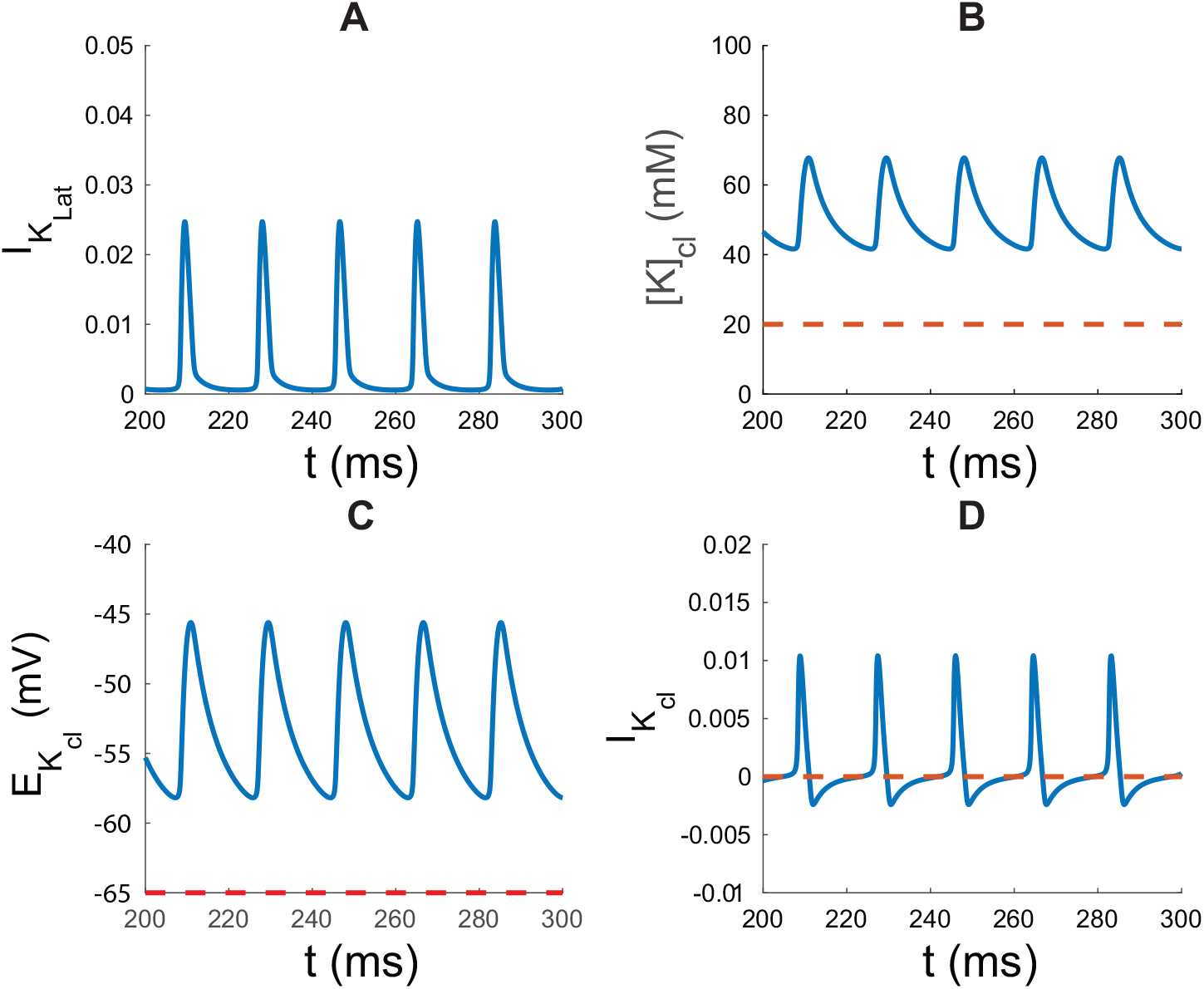
Panel A: Lateral potassium current 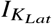, Panel B: Cleft potassium concentration [*K*]_*cl*_, Panel C: Junctional membrane Nernst potential 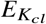, and Panel D: Junctional potassium current 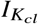 for fraction of potassium cleft current *F*_*I K*_ = 0.7, and nominal cleft width *w* _*f*_ = 1. (A) Potassium current 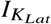 on the lateral membrane is always an outward current and rapidly changes during an action potential since extracellular potassium and reversal potential are invariant along the membrane facing bulk unrestricted volumes. (B) Extracellular potassium in the cleft ([*K*]_*cl*_) (blue curve) is elevated relative to invariant bulk extracellular potassium [*K*]_*e*_ = 20mM (dashed red line) during inter-action potential intervals. (C) The cleft potassium equilibrium potential 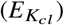 (blue curve) is elevated relative to bulk (red dashed line). During the inter-action potential interval 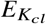 decreases early after repolarization and begins to rise as the lateral membrane begins to depolarize. (D) Cleft potassium current 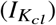 (blue curve) is inward (i.e.negative) early during the inter-action potential interval, becomes transiently outward (i.e., positive) as the cell begins to depolarize, strongly outward during the early phase of the action potential, and strongly inward for a prolonged period of time during repolarization.

Before further exploring the consequences of this cellular oscillation phenomenon, it is important to consider the following. Many of the potassium channels active during the diastolic potential in cardiomyocytes or inter-action potential of neurons exhibit strong inward rectification below the potassium reversal potential - a transmembrane potential below which an excitable cell should theoretically never fall. Interestingly, cellular and sub-cellular structural biologists are continuously exploring how protein cellular distributions, clustering, and nano-environments confer distinctly different emergent biophysical behaviors and local control on cells. Given the prevalence of strongly inward rectifying potassium channels in multiple cell types, (32)(33), all of which can exhibit some form of intrinsic organized activity, the model suggests that a relatively small, shared extracellular domain can easily coordinate synchronized and cyclic activity in tissue.

### Effects of Varying Fractional Cleft Potassium Current

In Figure 5 is shown the behavior of the model as a function of the cleft potassium current fraction *F*_*I K*_. Below the threshold of *F*_*I K*_ = 0.62, there are no spontaneous oscillations. Below this threshold value, the resting membrane potential is a gradually increasing function of *F*_*I K*_ (red curve -Panel A), as is the concentration of potassium in the cleft [*K*]_*cl*_ (red curve -Panel B). However, once fractional cleft potassium current *F*_*I K*_ exceeds 0.62, oscillatory activity begins. The oscillatory behavior in this parameter range is associated with the increased average level of potassium in the cleft [*K*]_*cl*_, similar to the behavior of the HH model. The oscillation of [*K*]_*cl*_ is a direct consequence of the oscillation of *V*_*m*_ which drives an oscillatory potassium current 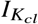 (Panel C). Interestingly, increasing *F*_*I K*_ in the range that supports spontaneous oscillatory activity can increase the frequency of spontaneous activity (Panel D). For this model and parameter space, the increased oscillatory frequency may be a result of the increasing [*K*_*cl*_] that increases the *m*-gate probability of activating sodium channels more than *h*-gate inactivation. However, as *F*_*I K*_ increases beyond 0.87, the average level of [*K*]_*cl*_ is too high to sustain oscillations, as is also the case in the HH model.

**Figure 5:**
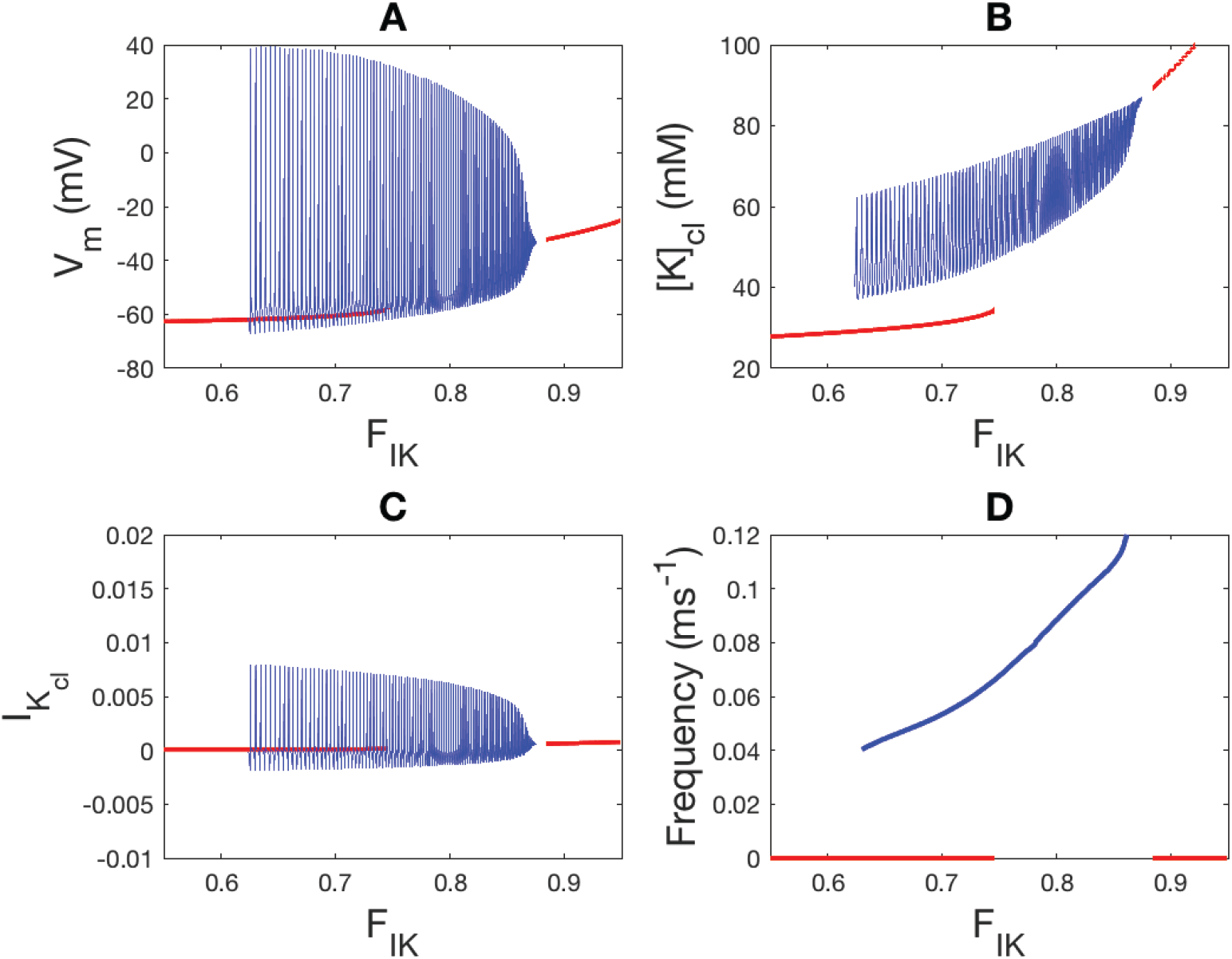
Solution diagram for the modified HH cleft model as a function of fractional potassium cleft current *F*_*I K*_ with cleft width fixed *w* _*f*_ = 1. Red curves represents stable steady state solutions and blue curves represent stable oscillatory solutions. Spontaneous oscillatory solutions exist for 0.62 < *F*_*I K*_ < 0.87.

Importantly, this model is highly dependent on the simplistic nature of *I*_*K*_, which does not account for the complex kinetics of the many inward rectifier potassium channel protein families, subtypes, and heteromeric combinations. Nonetheless, one expects that specialized sub-cellular localization, coupled with the unique biophysical channel properties will produce emergent oscillatory behaviors even in different cell types.

### Effects of Increasing Cleft Width

Based on what has been seen so far, we hypothesize that increasing cleft width should have the effect of decreasing the likelihood of spontaneous oscillations occurring. This is because with increased cleft width, the outward potassium cleft current has less influence on the potassium cleft concentration, and increased cleft width increases the cleft diffusion coefficient *δ*_*cl*_ and decreases the cleft resistance *r*_*cl*_, thus keeping cleft potassium concentration [*K*]_*cl*_ closer to the extracellular potassium concentration and the cleft potential *V*_*cl*_ closer to *V*_*e*_ = 0. This is what is observed in Figure 6. There we see the effect of varying *w* _*f*_ while keeping *F*_*I K*_ = 0.75 fixed. Specifically, at small values of *w* _*f*_, potassium accumulation in the cleft is substantial and has a large amplitude oscillation. (Panel B) As *w* _*f*_ increases, the amplitude of potassium concentration oscillation decreases as does the average potassium concentration level. For sufficiently large *w* _*f*_, (here *w* _*f*_ > 7.5), potassium accumulation is too small to drive oscillations and the oscillations cease to exist, and cleft concentration is not much different from the extracellular concentration.

**Figure 6:**
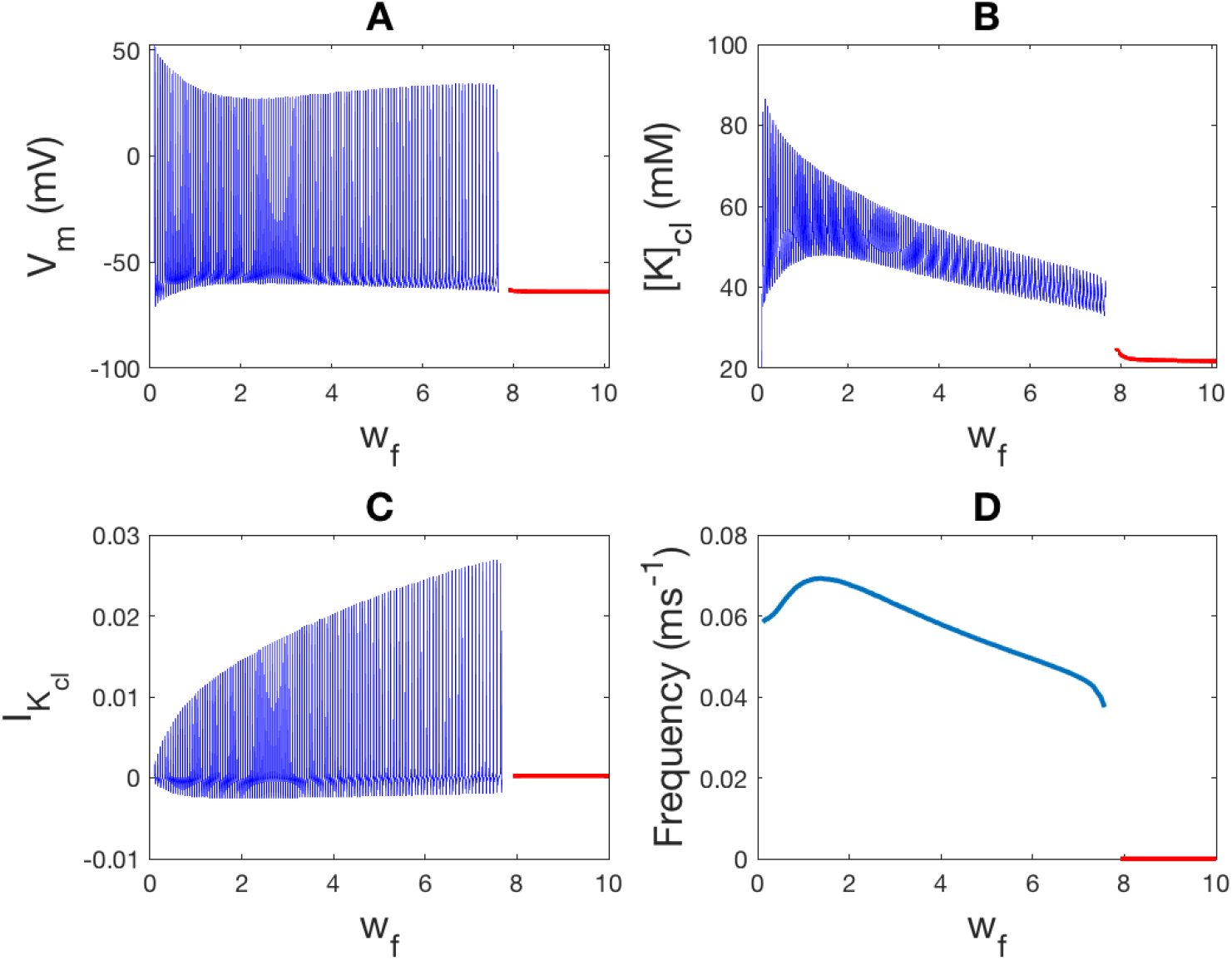
Solution diagram for the modified HH cleft model as a function of cleft width *w* _*f*_ with fixed fractional potassium cleft current *F*_*I K*_ = 0.75. Spontaneous oscillatory solutions (blue curves) disappear for *w* _*f*_ > 7.5. For *w* _*f*_ > 7.5 the solution (red curve) is at a stable steady state. The amplitude of potassium concentration oscillations and average value of potassium cleft concentration (Panel B) is a decreasing function of *w* _*f*_.

Interestingly, the amplitude of the oscillatory potassium current 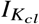 is increasing as *w* _*f*_ increases (Panel C), for the simple reason that the potassium Nernst potential is decreasing, hence increasing the amplitude of the potassium current driving force. Similar to the findings in Figure 5 Panel D, spontaneous oscillatory activity in Figure 6 Panel D is not constant but is dependent on *w* _*f*_. In this case, the oscillatory frequency can be complex and multiphasic, and appears to once again follow the diastolic [*K*]_*cl*_ values in Panel B. These data reveal that potassium regulation in narrow extracellular clefts can not only facilitate or repress spontaneous activity, but can also modulate the frequency of the spontaneous activity.

### Extracellular Coupled Oscillator Synchronization

Earlier, we posited that synchronized activity arising from spontaneous individual cellular activity is statistically unlikely without some kind of synchronizing process. Based on the ability of a single cell to oscillate in predictable patterns when a portion of its potassium channels face a restricted volume, it should come as no surprise that two cells sharing a restricted cleft will oscillate and entrain.

With parameter values *F*_*I K*_ = 0.59 and *w* _*f*_ = 1, the single cell model does not spontaneously oscillate. However, when *F*_*I K*_ = 0.59 for two cells sharing a restricted volume, they synchronously and simultaneously depolarize as expected (data not shown). Intercellular heterogeneity is a more interesting case, however. Figure 7 demonstrates that with *F*_*I K*_ = 0.65 in cell 1 and *F*_*I K*_ = 0.25 in cell 2 there is temporally coordinated, although not simultaneous, spontaneous oscillations. Specifically, the two cells concurrently increase [*K*]_*cl*_ in the shared cleft more rapidly than the individual cells alone, but there is a delay between action potential activations, demonstrating the synchronization can be the result of extracellular potassium mediated electrical excitation.

**Figure 7:**
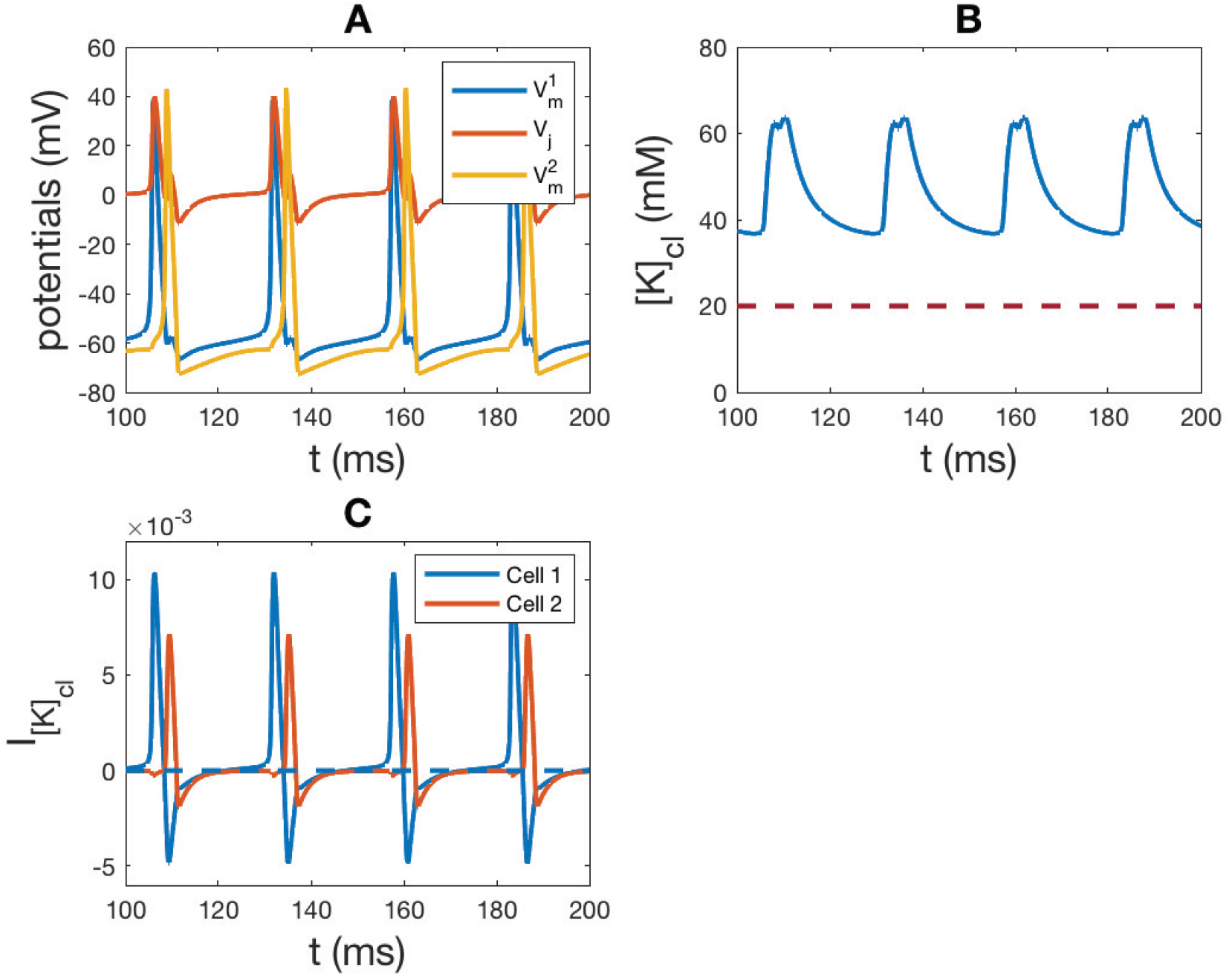
Panel A: Two cells sharing a restricted cleft spontaneously and synchronously activate even when fractional 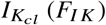 is heterogeneous. Here *F*_*I K*_ = 0.65 for cell 1 and *F*_*I K*_ = 0.25 for cell 2. Panel B: Electrophysiologic heterogeneity between cell 1 and 2 results in multiphasic potassium cycling resulting from temporally shifted [*K*]_*cl*_ dynamics. Panel B: The action potential of cell 1 increases [*K*]_*cl*_ (blue curve), and activation of cell 2 further increases [*K*]_*cl*_ until both currents are inward and in conjunction deplete [*K*]_*cl*_. Panel C: 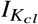 is different for the two apposing cell ends sharing the cleft. 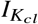 in cell 1 and 2 is biphasic during the respective action potentials, but the amplitude is reduced in cell 2. Co-dependence on cleft potassium is the mechanism that homogenizes action potential repolarization occurs without gap junctions.

Figure 7 Panel C reveals that 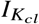 for cell 1 becomes positive after action potential activation and the shared [*K*]_*cl*_ rises dramatically (Figure 7 Panel B). This rapid rise in shared [*K*]_*cl*_ causes 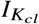 in cell 2 to transiently conduct inwardly and depolarize cell 2 by a potassium mediated pathway. The large inward and outward 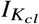 of cell 2 is less than 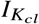 in cell 1, because of cleft priming caused by [*K*]_*cl*_ in cell 1 and the smaller number of potassium channels for cell 2. Interestingly, this inward 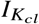 in cell 1 prolongs the late phase of the action potential in cell 1 (Figure 7 Panel A), suggesting that repolarization heterogeneity between electrically active cells can be reduced by a gap junction independent mechanism.

The model also suggests novel mechanistic roles for inexcitable cells (such as astrocytes, fibroblasts, and macrophages) co-distributed and interacting with excitable cells (e.g. neurons and myocytes). Astrocytes are known to modulate extracellular potassium and neuronal excitability in the tripartite synapse [40]. In the heart, fibroblasts are densely expressed in the pacemaking sinoatrial node (39)(40), and macrophages have been implicated as regulators of atrioventricular node conduction (41). While a previous study ruled out the role of capacitive coupling between cardiomyocytes and fibroblasts (39), it is important to note that capacitive coupling assumes excitation of the distal cell membrane without ion channels, and this is different from ephaptic coupling.

Representative action potentials in Figure 8 Panel A demonstrates what happens when an excitable cell (left) is coupled with an inexcitable cell (right). Specifically, cell 1 has currents of the original HH constants and kinetics, while Cell 2 is excitable with normal *g*_*N a*_ (Panel B, solid trace) or inexcitable with no sodium current *g*_*N a*_ = 0, (dashed trace). As expected, when the two cells are identical, *V*_*m*_ in both cells 1 and 2 oscillates (*F*_*I K*_ = 0.6). When cell 2 is inexcitable with *F*_*I K*_ = 0.6, there is no oscillation (data not shown). However, increasing *F*_*I K*_ in the inexcitable cell 2 (to *F*_*I K*_ = 0.9) reveals that the inexcitable cell induces spontaneous oscillations in the excitable cell 1, but with a reduced frequency (dashed trace, Panel A). It is noteworthy that the potassium current in cell 2 (- dashed trace in Figure 8 B) is inward, partially depleting potassium from the cleft, but the amplitude of the potassium current in cell 1 is substantially enhanced compared to the situation where the two cells are identical (solid trace). As a result, peak [*K*]_*cl*_ in Figure 8 C is only modestly reduced in the heterogeneous cell pair (dashed trace) relative to the homogeneous cell pair, because 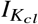 is enhanced into cell 1 despite negligent contributions of 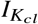 from cell 2. In summary, this model predicts that non-excitable cells can modulate excitability as previously proposed, but they can also modulate spontaneous activity and modify the frequency response of spontaneous activity to a range of heterogeneous anatomical and functional protein expression patterns, including preventing spontaneous activity altogether.

**Figure 8:**
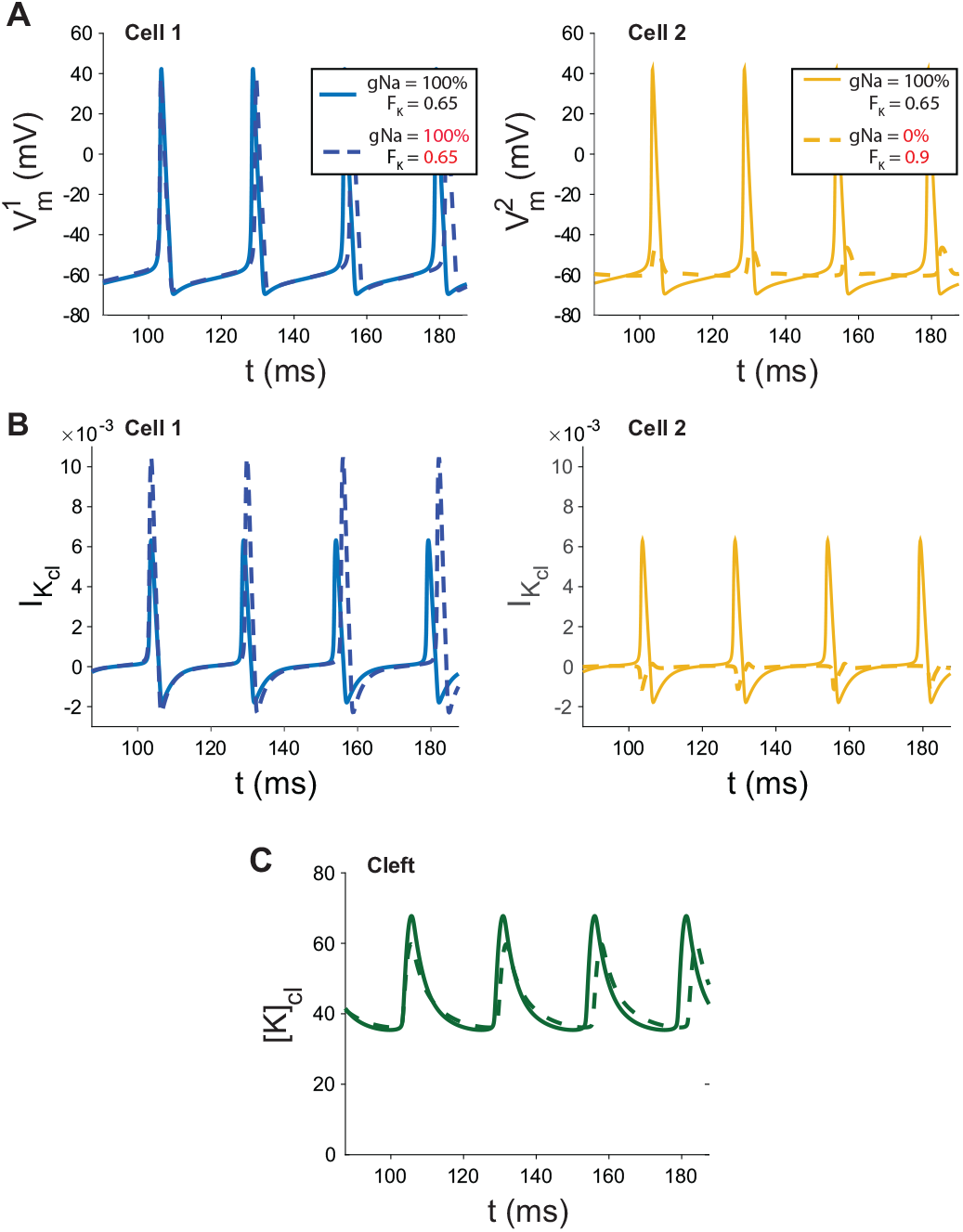
Panel A: Oscillatory solutions for the dual cell model with cells coupled via a common narrow cleft domain (*w* _*f*_ = 1) and fractional potassium cleft current *F*_*I K*_ = 0.65 for both cells, cell 2 with standard sodium channel density (solid curves) and no sodium channels (dashed curves) hence inexcitable. Excitable cell 1 spontaneously activates without significantly affecting the transmembrane potential *V*_*m*_ of inexcitable cell 2. Spontaneous activity frequency is decreased in excitable cell 1 when fractional cleft 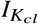 of inexcitable cell 2 is increased (dashed lines) because of cleft potassium withdrawal by cell 2 causing a small transmembrane depolarization that follows activation of cell 1. Panel B: Cleft potassium currents corresponding to the potential oscillations *V*_*m*_ shown in A, with both cells identical and excitable (solid curves), and cell 1 excitable with *F*_*I K*_ = 0.65 (Panel A - dashed) and cell 2 inexcitable with *F*_*I K*_ = 0.9 (Panel B - dashed). Panel C: Peak [*K*]_*cl*_ is only partially reduced in the electrophysiologically heterogeneous cell pairs because the outward 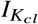 is partially counter balanced by the small inward 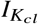 into cell 2.

## DISCUSSION

### Summary and Conclusions

The addition of an extracellular volume with limited potassium diffusivity can cause spontaneous oscillatory behavior in an otherwise quiescent in silico cell model by a mechanism where potassium channels can inwardly rectify in the cleft to depolarize cells. The model also suggests that synchronized oscillation between multiple cells can occur in the absence of gap junctional coupling. Importantly, shared extracellular volumes may be a new mechanism to reduce dispersion of repolarization, also by a gap junction independent mechanism. Interestingly, the restricted extracellular volume clock hypothesis may offer some insights into why some groups report that cultured cells do and do not spontaneously and regularly fire action potentials (10)(12)(43). Most cells in culture are adhered to surfaces like plastics and glass with proteins of different lengths. One could even hypothesize that choice of adhesion protein like laminin or fibronectin, for examples, will affect automaticity. While a retrospective review of the literature would support this hypothesis, experimental studies should be performed to estimate the resistance between the plate and cells in order to determine if the cell-surface interface provides a sufficient restricted volume for a substantial portion of sarcolemmal ion channels to facilitate spontaneous activity.

The model also suggests a novel role for excitable cells structurally coupled to inexcitable cells. For example, there is still active debate in the cardiac field on whether the relatively small volume of the inexcitable fibroblast substantially alters resting membrane potential of cardiomyocytes if electrically coupled by gap junctions. This work suggests that a fibroblast may also modulate excitable cells in the same way an astrocyte spatially buffers extracellular ions and neurotransmitters in the tripartite synapse (44). With regards to neuroscience, the models suggest that support cell types like astrocytes may switch from suppressing to facilitating spontaneous neuronal activity in synaptic groups if, for examples, outward potassium currents are reduced in neurons or enhanced in astrocytes. More generally, the data demonstrate the possibility that shared extracellular nano-environments create emergent phenomenon that cannot be solely accounted for by trans and intracellular pathways. Lastly, in accordance with the “inverted cable” concept of ephaptic conduction proposed in (38), it may be advantageous to modify our predominant frame that cellular excitability refers to transmembrane responses considered mainly from an intracellular perspective, and consider that inexcitable cells may have inverted action potentials.

## AUTHOR CONTRIBUTIONS

SP conceptualized the hypothesis and initial model, and co-wrote the manuscript. JPP developed the hypothesis, a physically accurate model and bifurcation analysis, and he co-wrote the manuscript

## ACKNOWLEDGMENTS

This study was supported by funding from the National Institutes of Health, grant numbers R01HL141855 (SP) R01HL102298 (SP, JPK) and R01HL138003 (SP).

## REFERENCES

1. DiFrancesco, D., Pacemaker mechanisms in cardiac tissue. Annu Rev Physiol, 1993. 55: p. 455–72.

2. Pape, H.C., Queer current and pacemaker: the hyperpolarization-activated cation current in neurons. Annu Rev Physiol, 1996. 58: p. 299–327.

3. Mitsuiye, T., Y. Shinagawa, and A. Noma, Sustained inward current during pacemaker depolarization in mammalian sinoatrial node cells. Circ Res, 2000. 87(2): p. 88–91.

4. Biel, M., A. Schneider, and C. Wahl, Cardiac HCN channels: structure, function, and modulation. Trends Cardiovasc Med, 2002. 12(5): p. 206–12.

5. Wilders, R., Computer modelling of the sinoatrial node. Med Biol Eng Comput, 2007. 45(2): p. 189–207.

6. Lakatta, E.G., V.A. Maltsev, and T.M. Vinogradova, A coupled SYSTEM of intracellular Ca2+ clocks and surface membrane voltage clocks controls the timekeeping mechanism of the heart’s pacemaker. Circ Res, 2010. 106(4): p. 659–73.

7. Irisawa, H., H.F. Brown, and W. Giles, Cardiac pacemaking in the sinoatrial node. Physiol Rev, 1993. 73(1): p. 197–227.

8. McCormick, D.A. and T. Bal, Sleep and arousal: thalamocortical mechanisms. Annu Rev Neurosci, 1997. 20: p. 185–215.

9. Ramirez, J.M. and D.W. Richter, The neuronal mechanisms of respiratory rhythm generation. Curr Opin Neurobiol, 1996. 6(6): p. 817–25.

10. Raman, I.M. and B.P. Bean, Ionic currents underlying spontaneous action potentials in isolated cerebellar Purkinje neurons. J Neurosci, 1999. 19(5): p. 1663–74.

11. Levic, S., P. Lv, and E.N. Yamoah, The activity of spontaneous action potentials in developing hair cells is regulated by Ca(2+)-dependence of a transient K+ current. PLoS One, 2011. 6(12): p. e29005.

12. Guo, X., et al., In vitro Differentiation of Functional Human Skeletal Myotubes in a Defined System. Biomater Sci, 2014. 2(1): p. 131–138.

13. Arrowsmith, S. et al., What do we know about what happens to myometrial function as women age? J Musc Res and Cell Mot. 2012. 33(3-4): p 209–17

14. Xie, Y., et al., So little source, so much sink: requirements for afterdepolarizations to propagate in tissue. Biophys J, 2010. 99(5): p. 1408–15.

15. Hodgkin, A.L. and A.F. Huxley, A quantitative description of membrane current and its application to conduction and excitation in nerve. J Physiol, 1952. 117(4): p. 500–44.

16. Melnyk, P., et al., Differential distribution of Kir2.1 and Kir2.3 subunits in canine atrium and ventricle. Am J Physiol Heart Circ Physiol, 2002. 283(3): p. H1123–33.

17. Gillet, L., et al., Cardiac-specific ablation of synapse-associated protein SAP97 in mice decreases potassium currents but not sodium current. Heart Rhythm, 2015. 12(1): p. 181–92.

18. Petitprez, S., et al., SAP97 and dystrophin macromolecular complexes determine two pools of cardiac sodium channels Nav1.5 in cardiomyocytes. Circ Res, 2011. 108(3): p. 294–304.

19. Milstein, M.L., et al., Dynamic reciprocity of sodium and potassium channel expression in a macromolecular complex controls cardiac excitability and arrhythmia. Proc Natl Acad Sci U S A, 2012. 109(31): p. E2134–43.

20. Matamoros, M., et al., Nav1.5 N-terminal domain binding to alpha1-syntrophin increases membrane density of human Kir2.1, Kir2.2 and Nav1.5 channels. Cardiovasc Res, 2016. 110(2): p. 279–90.

21. Eichel, C.A., et al., Lateral Membrane-Specific MAGUK CASK Down-Regulates NaV1.5 Channel in Cardiac Myocytes. Circ Res, 2016. 119(4): p. 544–56.

22. Shy, D., et al., PDZ domain-binding motif regulates cardiomyocyte compartment-specific NaV1.5 channel expression and function. Circulation, 2014. 130(2): p. 147–60.

23. Veeraraghavan, R., et al., The adhesion function of the sodium channel beta subunit (beta1) contributes to cardiac action potential propagation. Elife, 2018. Aug 14;7.

24. Veeraraghavan, R., et al., Sodium channels in the Cx43 gap junction perinexus may constitute a cardiac ephapse: an experimental and modeling study. Pflugers Arch, 2015. 467(10): p. 2093–105.

25. Veeraraghavan, R., et al., Potassium channels in the Cx43 gap junction perinexus modulate ephaptic coupling: an experimental and modeling study. Pflugers Arch, 2016. 468(10): p. 1651–61.

26. Ponce-Balbuena, D., et al., Cardiac Kir2.1 and NaV1.5 Channels Traffic Together to the Sarcolemma to Control Excitability. Circ Res, 2018. 122(11): p. 1501–1516.

27. Rhett, J.M., J. Jourdan, and R.G. Gourdie, Connexin 43 connexon to gap junction transition is regulated by zonula occludens-1. Mol Biol Cell, 2011. 22(9): p. 1516–28.

28. Greer-Short, A., et al., Revealing the Concealed Nature of Long-QT Type 3 Syndrome. Circ Arrhythm Electrophysiol, 2017. 10(2): p. e004400.

29. Novak, M., et al., Intercellular sodium regulates repolarization in cardiac tissue with sodium channel gain-of-function. Biophysical J, 2020. IN PRESS

30. Ashley, L.M., A determination of the diameters of ventricular myocardial fibers in man and other mammals. Am J Anat, 1945. 77: p. 325–63.

31. Raisch, T., M. Khan, and S. Poelzing, Quantifying Intermembrane Distances with Serial Image Dilations. J Vis Exp, 2018(139).

32. Ransom, C.B. and H. Sontheimer, Biophysical and pharmacological characterization of inwardly rectifying K+ currents in rat spinal cord astrocytes. J Neurophysiol, 1995. 73(1): p. 333–46.

33. Hibino, H., et al., Inwardly rectifying potassium channels: their structure, function, and physiological roles. Physiol Rev, 2010. 90(1): p. 291–366.

34. Kucera, J.P., S. Rohr, and Y. Rudy, Localization of sodium channels in intercalated discs modulates cardiac conduction. Circ Res, 2002. 91(12): p. 1176–82.

35. Sperelakis, N., Combined electric field and gap junctions on propagation of action potentials in cardiac muscle and smooth muscle in PSpice simulation. J Electrocardiol, 2003. 36(4): p. 279–93.

36. Mori, Y., G.I. Fishman, and C.S. Peskin, Ephaptic conduction in a cardiac strand model with 3D electrodiffusion. Proc Natl Acad Sci U S A, 2008. 105(17): p. 6463–8.

37. Lin, J. and J.P. Keener, Modeling electrical activity of myocardial cells incorporating the effects of ephaptic coupling. Proc Natl Acad Sci U S A, 2010. 107(49): p. 20935–40.

38. Lin, J. and J.P. Keener, Ephaptic coupling in cardiac myocytes. IEEE Trans Biomed Eng, 2013. 60(2): p. 576–82.

39. De Maziere, A.M., et al., Spatial and functional relationship between myocytes and fibroblasts in the rabbit sinoatrial node. J Mol Cell Cardiol, 1992. 24(6): p. 567–78.

40. Kohl, P., et al., Mechanosensitive fibroblasts in the sino-atrial node region of rat heart: interaction with cardiomyocytes and possible role. Exp Physiol, 1994. 79(6): p. 943–56.

41. Hulsmans, M., et al., Macrophages Facilitate Electrical Conduction in the Heart. Cell, 2017. 169(3): p. 510–522 e20.

42. Nwaobi, S.E., et al., The role of glial-specific Kir4.1 in normal and pathological states of the CNS. Acta Neuropathol, 2016. 132(1): p. 1–21.

43. Grubic, Z., et al., Myoblast fusion and innervation with rat motor nerve alter distribution of acetylcholinesterase and its mRNA in cultures of human muscle. Neuron, 1995. 14(2): p. 317–27.

44. Bolton, S., et al., Regulation of the astrocyte resting membrane potential by cyclic AMP and protein kinase A. Glia, 2006. 54(4): p. 316–28.

